# Neural and mechanical properties of vastus lateralis and vastus medialis at different rectus femoris muscle lengths

**DOI:** 10.64898/2026.02.03.703444

**Authors:** Milena A. dos Santos, Hélio V. Cabral, J Greig Inglis, Caterina Cosentino, Elmira Pourreza, Liliam Fernandes de Oliveira, Francesco Negro

**Author notes:** Corresponding author: Prof. Francesco Negro, Department of Clinical and Experimental Sciences, Università degli Studi di Brescia, Viale Europa 11, Brescia, 25121, Italy.

## Abstract

This study investigated how altering the length of one muscle influences motor unit discharge behavior of its synergists. Eighteen participants performed isometric knee extensions at 10% and 30% of maximal voluntary contraction (MVC) with the hip joint at 90° (shortened rectus femoris, RF) and 180° (lengthened RF). High-density surface electromyograms were recorded from vastus medialis, RF and vastus lateralis, and decomposed into motor unit spike trains. Mean discharge rate and coefficient of variation of interspike interval were analyzed for tracked units in the vasti and non-tracked in the RF. While no changes were observed in RF motor units, lengthening the RF led to increased discharge rates of vasti motor units at 10% MVC, but not 30% MVC. To further explore these force-dependent changes, two further experiments were conducted. The first showed that the discharge rate at recruitment during ramp-up contractions increased with RF lengthening, but only for low-threshold vasti units. In the second, electrically evoked twitch contractions in the vasti revealed significantly reduced twitches at 180° during low-frequency, but not high-frequency stimulation. These findings collectively suggest that the force-dependent changes in the vasti motor unit discharge rates are likely driven by RF-length dependent changes in the vasti muscles’ contractile properties.

## Introduction

During voluntary muscle contraction, alpha motoneurons integrate synaptic inputs from cortical and spinal sources to generate the neural drive to the muscle [1] and are therefore considered the final common pathway of the neuromuscular system [2]. Consequently, the force exerted by a muscle directly depends on the central and peripheral properties of motor units (i.e., the alpha motoneurons and the muscle fibers they innervate) [3,4]. Across most of a muscle’s operating range, changes in force output are achieved through adjustments in the number of active motor units (recruitment) and their discharge rates (rate coding), mechanisms that primarily reflect central neural control properties [3,5]. The relative contribution of recruitment and rate coding across force levels has been shown to be muscle- and task-dependent and is influenced by several factors, including the functional role of the muscle [6]. In parallel, peripheral motor unit properties are tightly linked to muscle-tendon unit mechanics, such as variations in conduction velocity and fiber length, which directly influence force generating capacity [7–9]. These neural and mechanical factors interact dynamically during contraction to ultimately determine force control [5,10,11].

The mechanical properties of the muscle-tendon unit arise from the interaction between active contractile elements and passive elastic structures, both within and external to the sarcomeres [12]. Active elements directly dictate force generation through ATP hydrolysis during actin-myosin cross-bridge cycling [13]. Passive structures within the sarcomere, such as titin, components of the myosin neck, and other structural proteins, also contribute to force production through their elastic properties [12,14]. Additional passive elements, including fasciae and tendons, also develop tension when elongated and exhibit mechanical properties adapted to their functional demands [15,16]. Because both active and passive components are strongly length-dependent, muscle force production varies substantially with changes in muscle length [17–19]. At shorter or longer lengths, active force depends on the degree of actin-myosin overlap, whereas passive structures store and release elastic energy to facilitate force transmission [12,20]. Thus, manipulating muscle length provides a unique framework for probing the interaction between central neural control and peripheral mechanical properties during force production [21,22].

Evidence from electrically evoked contractions has demonstrated that higher stimulation frequencies are required to achieve similar force outputs at shorter muscle lengths than at longer lengths, suggesting compensatory increases in motor unit discharge rates under mechanically disadvantaged conditions [23–26]. In contrast, findings from voluntary contractions are more variable. Some studies have reported that changes in muscle-tendon length induced by joint position do not significantly affect motor unit discharge rate [27–29], whereas others have observed increased recruitment thresholds at shorter muscle lengths [30–32]. Moreover, alterations in muscle length during voluntary contractions influence other aspects of neural control, such as Ia afferent feedback, which modulates force variability [33]. Although this body of research has provided important insights into the neuromechanical interplay underlying force control, most studies have focused on monoarticular muscles and examined the effect of length-dependent changes on their own motor unit control. Much less is known about how changes in the mechanical properties of one muscle influence the neural control of motor units in its synergists. This question is particularly relevant because synergistic muscles, while modularly coordinated by the central nervous system [34,35] and sharing a substantial portion of their synaptic inputs [36,37], may present distinct lengths depending on joint configuration and task demands.

Therefore, the aim of this study was to investigate how changes in the length of one muscle influence the motor unit behavior of its synergistic muscles. The quadriceps muscle group provides an ideal model for this investigation, as knee extension force is generated by both monoarticular muscles acting exclusively on the knee joint (vastus lateralis (VL), vastus medialis (VM) and vastus intermedius) and the biarticular muscle rectus femoris (RF), which spans both the knee and hip joints [38]. By altering the hip joint angle from 90° (seated position) to 180° (supine position), we selectively modified RF muscle length without affecting the length of monoarticular vasti, allowing us to isolate the effect of RF length changes on the discharge behavior of VM and VL motor units. To support the interpretation of results of the primary experiment, two additional experiments were performed. In the first, participants performed ramp-up isometric knee extension contractions at both RF muscle lengths to examine force-dependent changes in motor unit discharge behavior. In the second, electrically evoked twitch contractions were elicited from the vasti muscles while manipulating RF length to determine whether neural adaptations in the vasti associated with RF length changes were accompanied by changes in their contractile properties. We hypothesize that lengthening the RF muscle would alter both central and peripheral motor unit properties in the VM and VL muscles.

## Methods

### Participants

Eighteen healthy individuals (10 males and 8 females, age 28 ± 6.4 years, mass 67.45 ± 19.09 kg and height 1.74 ± 0.1 m) volunteered to participate in this study. All participants had no history of lower limb injury or pain that could impact on their ability to perform voluntary contractions during the experiments. Prior to participating in the experiments, all subjects provided written informed consent. This study was conducted in accordance with the latest version of the Declaration of Helsinki and approved by the local ethics committee of the University of Brescia (code 5665).

### Experimental protocol

#### Experiment 1

Participants were seated on the isokinetic dynamometer (HUMACNORM Extremity System, CSMi Solutions, Stoughton, MA, USA) with their right knee flexed at 90° (0° = full extension) and the lateral femoral condyle aligned with the axis of rotation of the dynamometer. The right ankle was secured to the dynamometer with straps approximately two fingers above the lateral malleolus to measure isometric knee extension torque. Participants were asked to keep their left leg relaxed during experiments. The protocol was repeated for two hip positions: 90° (seated) and 180° (supine) (**Figure 1A**). During the transition from 90° to 180°, the rectus femoris was lengthened.

**Figure 1.**
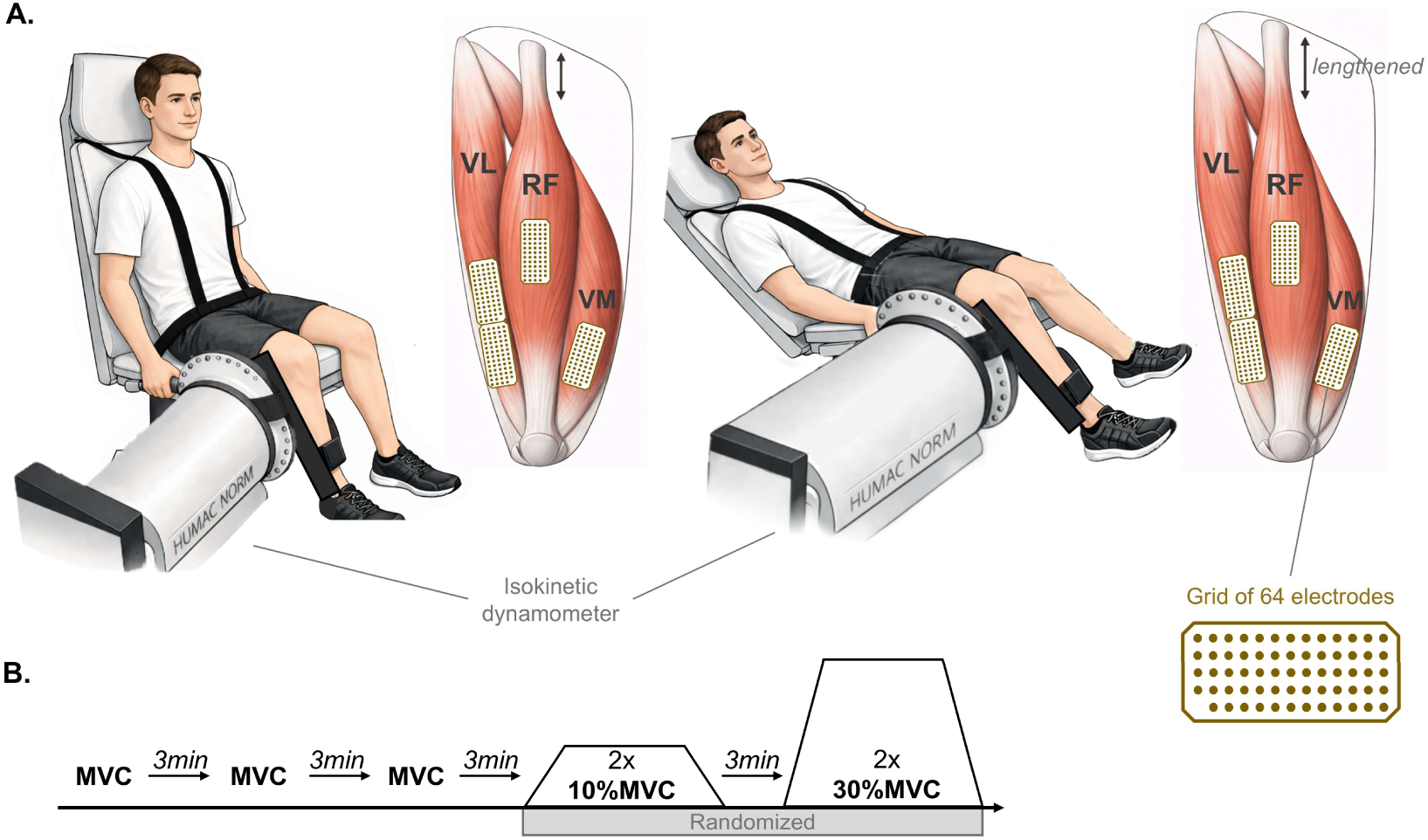
(A) Schematic representation of the experimental setup for measuring knee extension isometric force with the hip positioned at 90° (rectus femoris was shortened; left panel) and at 180° (rectus femoris was lengthened; right panel. High-density surface electromyograms were acquired from the vastus medialis (VM), rectus femoris (RF), and vastus lateralis (VL) muscles using grids of 64 electrodes. (B) Experimental tasks performed by the participants. After maximal voluntary contractions (MVC) of knee extension, participants were asked to follow trapezoidal force profiles, both at 10% MVC and 30% MVC. The image of the participant in panel A was generated by the authors (MAS, HVC) in PowerPoint (Office 365, Microsoft) and enhanced using ChatGPT (v. 5.5, https://chatgpt.com/).

Participants first completed three isometric maximal voluntary contractions (MVC) for 3 s, with 3 min of rest between trials. The highest peak torque from the three MVCs was considered as a reference for the following submaximal isometric contractions. After 3 min rest, participants performed isometric torque-matching tasks following a trapezoidal profile. The trapezoidal task consisted of increasing the isometric knee extension torque from 0% MVC to the target torque of 10% or 30% of MVC/s, maintaining the target torque for 30 s (plateau phase), and returning from the target torque to 0% MVC at 10% MVC/s. The trapezoidal tasks were repeated twice for each target force, in a randomized order. The detailed experimental protocol is schematically represented in **Figure 1B**. Both the target and produced torques were displayed on a monitor using a custom-written MATLAB (version R2022b, The Mathworks Inc., Natick, MA, USA) interface.

To verify that altering the hip position would cause changes only to the RF muscle, B-mode ultrasound imaging (Telemed Medical Systems, Echowave II (x64) v 4.3.0, Milan, Italy, ArtUS) was acquired from a subgroup of 11 participants (2 female) during rest. A skin marking pencil was used to draw a reference line from the anterior superior iliac spine to the superior border of the patella. From this line, two others were traced over the skin at 20° laterally and 45° medially. The three references lines were used to guide the researcher on finding the RF, VL and VM muscles, respectively[39] (**Figure 2A**). For both supine and seated positions, the proximal and distal musculotendinous junctions of the three muscles were identified by the same researcher with the ultrasound probe positioned longitudinally to the muscle. **Figure 2B** shows examples of ultrasound images of the distal part of the three muscles, with numbers indicating the distal musculotendinous junctions marked on the skin in **Figure 2A**. Each musculotendinous junction was verified at least three times before establishing the final mark. A 64 mm linear probe operating at 9 MHz (LF9-5N60-A3, penetration depth range of 40-65mm) was used. Then, with the aid of a measuring tape similar to a previous study[40], the length of the three muscles was estimated from the distance between the proximal and distal musculotendinous junctions marked on the skin and normalized to the estimated femur length. The estimated femur length was calculated from the greater trochanter to the lateral condyle of the femur. All measurements were first performed with the participants on supine position. Before changing hip positions, the pencil marks were cleaned to reduce bias.

**Figure 2.**
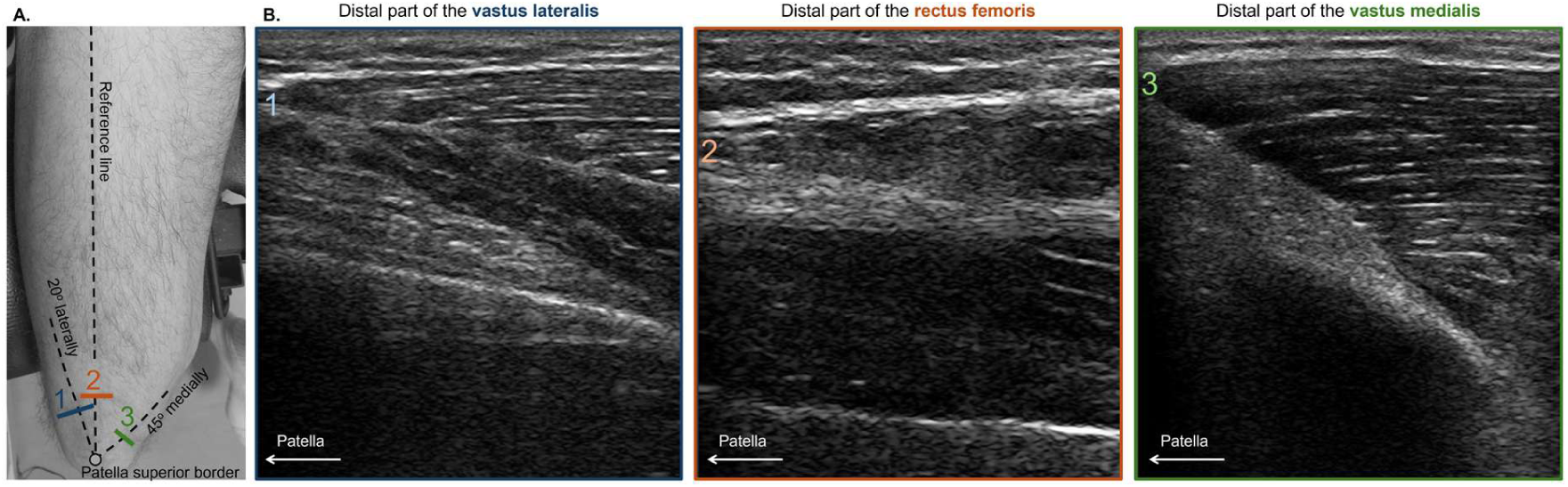
Schematic representation of the steps involved during the ultrasound imaging of the musculotendinous junction from VL, RF and VM. (A) Photo of the tight with the reference lines used to guide the evaluator followed by respective marks and numbers 1, 2 and 3 indicating the VL, RF and VM distal musculotendinous junctions and (B) its corresponding ultrasound image.

To investigate the observed force-dependent changes in the vasti motor unit discharge rate in this experiment (see *Results*), we conducted two further sets of experiments (*Experiment 2 and Experiment 3*).

#### Experiment 2

Experiment 2 investigated whether observed force-dependent changes were related to the motor unit recruitment thresholds, as different thresholds of motor units have been shown to present different twitch properties. For this, the same participants from Experiment 1 performed ramp-up isometric knee extension contractions, reaching 30% MVC at a rate of 10% MVC/s. The task was repeated twice for both hip positions (90° and 180°). Participants were positioned in the dynamometer chair as described in *Experiment 1*.

#### Experiment 3

Experiment 3 directly explored the effect of changing the RF on the vasti twitch profiles in a subgroup of seven healthy individuals (4 females; age 28.4 ± 4.46 years, weight 69.71 ± 17.73 kg and height 1.71 ± 0.95 m; three participants in common with the previous experiment). In this experiment, electrical stimulation was applied over the VL and VM muscles while participants were positioned as described in the voluntary contraction experiments (**Figure 1A**). Initially, an experienced researcher identified the VL and VM dominant motor points as described previously [41]. Briefly, after familiarizing the participants with the electrical stimulation sensation, the motor point identified using a pen electrode (cathode; Digitimer, Welwyn Garden City, United Kingdon) positioned over the distal region of the muscle and a rectangular anode electrode (size 45 x 80 mm, FIAB, Florence, Italy) fixed on the opposite side just above the popliteal fossa. The position of the cathode electrode was then displaced slightly until the location which required the least amount of stimulus intensity to elicit a muscle twitch was identified. This was considered the location of the dominant motor point. For each cathode position a train of 10-15 monophasic rectangular pulses at 1 Hz was delivered, and this procedure was performed separately for each muscle. After the motor point was identified bipolar derivation were implemented where a square cathode adhesive electrode (size 46 x 46 mm) was positioned over the motor point, and a rectangular anode electrode (size 45 x 80 mm) was placed over the muscle approximately 10 cm above the cathode. This electrode placement was based on piloting data and adapted from previous research [42,43]. For each hip position (90° and 180°), current pulses were delivered simultaneously over the VM and VL using two different stimulator devices (DS5 Isolated Bipolar Constant Current Stimulator, Digitimer, Welwyn Garden City, United Kingdon). Specifically, two 10-s trains of biphasic square pulses (1ms duration) were delivered, one at 10 Hz and the other at 20 Hz. These two frequencies included a variability of 80% for each muscle. The absence of frequency periodicity was used to obtain a unique solution when applying the deconvolution (see *Data analysis*) [44]. The amplitude of stimulation was set at 15 mA. The amplitude and stimulation frequencies were chosen based on piloting data with the aim of resembling the first experiment, where a lower-level torque (lower stimulation frequency) and a higher-level torque (higher stimulation frequency) were used.

### Data collection

High-density surface electromyograms (HDsEMG) were collected during submaximal voluntary contractions using four 64-electrode grids arranged in 13 rows by 5 columns each, with one missing electrode in the one corner (interelectrode distance of 8 mm; GR08MM1305, OT Bioelettronica, Turin, IT). The grids were placed over the distal region of the VM muscle, the central region of the RF and the proximal and distal regions of the VL muscle (**Figure 1C**), which were localized via palpation. Because the number of decomposed VM motor units was considerably low (see *Results*), for the last 7 subjects three grids of 64 electrodes with smaller interelectrode distance were placed over distal part of the VM (interelectrode distance of 4 mm; GR04MM1305, OT Bioelettronica, Turin, IT). This choice was based on recent evidence suggesting that a larger number of electrodes with a smaller interelectrode distance would lead to a greater motor unit yield [45].

Prior to electrode placement, muscle regions were shaved, mildly abraded (EVERI, Spes Medica), and cleaned with water. Additionally, an adhesive foam was filled with conductive paste (AC cream, Spes Medica) was applied to improve electrode-skin contact and reduce electrode movement. Two reference electrodes were utilized: one positioned over the right patella and another around the right wrist. HDsEMG signals were acquired in monopolar derivation, amplified by a factor of 150, and digitized at a sampling frequency of 2048 Hz using a 16-bit analog-to-digital converter (10-500 Hz bandwidth; Quattrocento, OT Bioelettronica, Turin, IT). Raw torque signals were sampled synchronously with HDsEMG signals.

### Data analysis

#### HDsEMG decomposition

The trial in which the participants followed the target force level more accurately (i.e., the lowest root-mean-square error between the target and the produced torque) was selected for further analysis. For the trapezoidal contractions, the analysis was performed on the central 30-s torque region (plateau). For the ramp-up task, the entire contraction was analyzed. First, HDsEMG signals were visually inspected, and bad channels, such as dead or those containing artifacts, were discarded. After visual inspection, the myoelectric signals recorded by the HDsEMG system from each electrode grid were decomposed into their individual motor unit spike trains using a convolutive blind source separation[46]. Only motor units with a silhouette value of 0.87, which is a measure of decomposition accuracy based on the separability between the identified discharge times and the baseline noise, were considered for further analysis. All decomposed motor units were visually inspected and any missed or misidentified motor unit discharges or discharge times, resulting in non-physiological discharge rates (i.e., < 4 Hz or > 50 Hz [47]), were manually edited by an experienced operator. This procedure has shown itself to be highly reliable across operators [48]. For the cases where we had more than one electrode grid on the same muscle, we removed the motor units that were identified more than once across grids before proceeding to tracking them between hip positions.

#### Motor unit tracking

To compare motor unit discharge behavior within the same motor units across hip positions, we tracked VM and VL motor units across hip angles in both Experiment 1 and Experiment 2 (**Figure 3**). We achieved this by reapplying the motor-unit-separation vectors estimated by the convolutive blind-source-separation algorithm from one hip position to the other, and vice versa. Subsequently, motor unit discharge rates were visually inspected and re-cleaned (when necessary). Due to changes in muscle length, the RF shifted relative to the surface electrode grid, preventing reliable tracking of motor units across hip positions. Therefore, non-matched RF motor units were used for this muscle for subsequent analyses.

**Figure 3.**
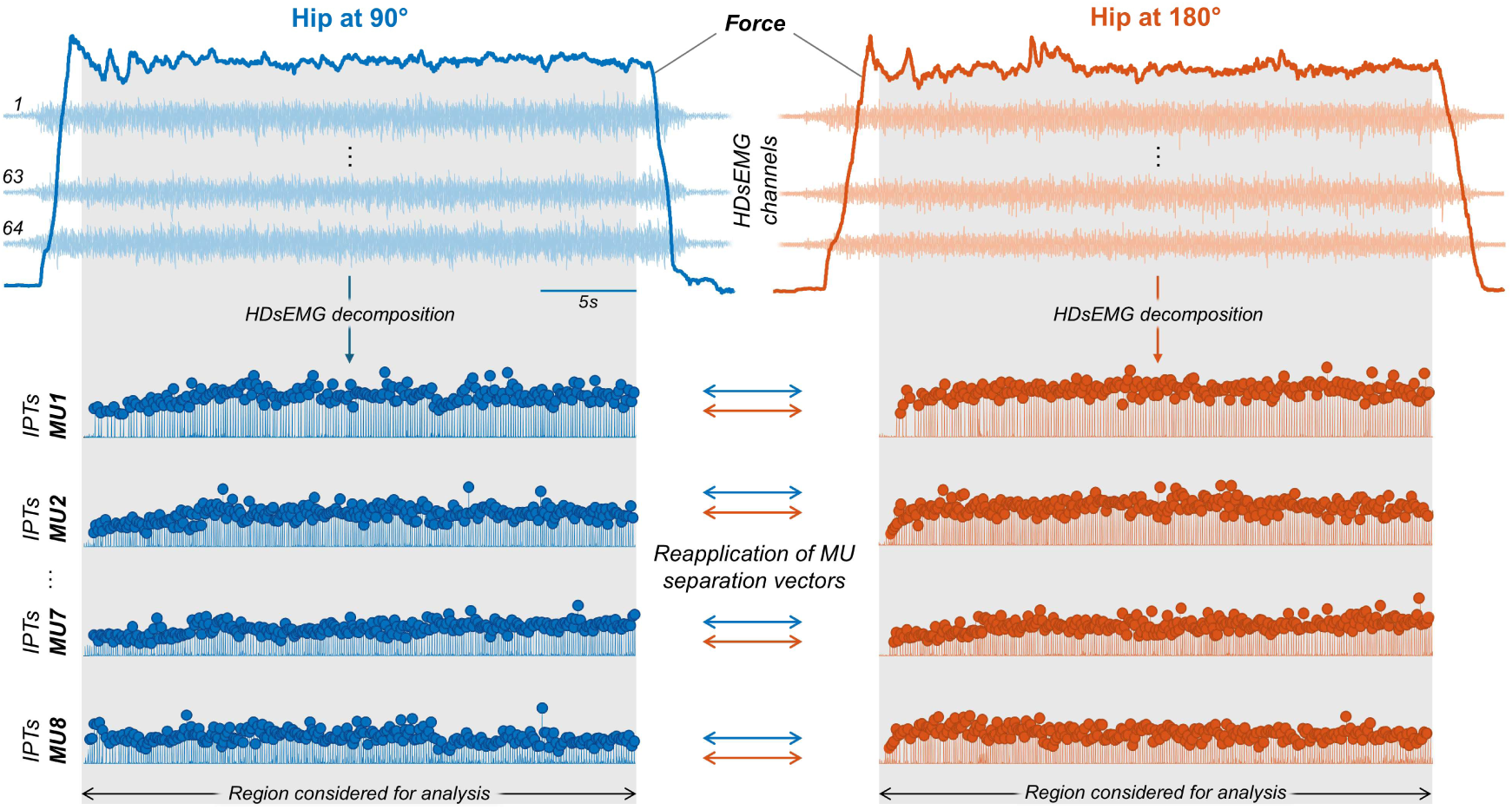
Steps involved in the motor unit data processing for trapezoidal and triangular contractions with the hip positioned at 90° (rectus femoris was shortened; blue) and at 180° (rectus femoris was lengthened; orange). The grey background indicates the time interval considered for analysis (plateau region). HDsEMG signals acquired using 64-electrodes grid were decomposed into individual motor unit spike trains. The bottom part shows examples of four innervation pulse trains of decomposed motor units, where each circle indicates the discharge time. The separation vectors of motor units decomposed at 90° were reapplied to the HDsEMG signals at 180°, and vice-versa, to track the same motor units across hip positions.

#### Motor unit discharge behavior

For the trapezoidal contractions (*Experiment 1*), the mean discharge rate and the coefficient of variation of interspike interval (CoV-ISI) of the matched VM and VL motor units, and non-matched RF motor units, were calculated. For the ramp up contractions (*Experiment 2*), we calculated the motor unit recruitment threshold force, the discharge rate at recruitment, and the mean discharge rate. For these variables, we divided the motor units into two groups: motor units recruited before 20% MVC and at or after 20% MVC. The first discharge was utilized to calculate the recruitment threshold [49]. The average of the first five discharges was used for the discharge rate at recruitment [50].

#### Torque analysis

For the trapezoidal task, the torque during the plateau was low-pass filtered at 15 Hz using a third-order Butterworth filter, and the coefficient of variation of torque was calculated to evaluate torque steadiness. For the evoked contractions, we estimated the average twitch profile representing the combined contribution of the VL and VM muscles to the total evoked torque. For this, a deconvolution method previously used to estimate individual motor unit force twitch profiles from discharge times of motor units and torque output during voluntary contractions was used [44]. The sum of the stimulation pulse trains applied to the VM and VL muscles was used to define the discharge times (i.e., stimulation times). Using these stimulation times, the combined VM-VL twitch profile, which, when convolved with the stimulation pulse trains, would reconstruct the recorded evoked torque were estimated. The variability in the respective stimulation times provided the possibility to invert the convolutive problem using a constrained least squares approach (MATLAB’s *lsqlin* function) performed separately for each hip position[51]. The peak twitch was calculated for each hip position and the proportion of change in percentage (from hip at 90° to hip at 180°) was retained for further analysis, separately for each frequency of stimulation. In addition, the element-wise average twitch across subjects was computed for each hip position and frequency of stimulation.

### Statistical analysis

All statistical analysis was performed in R Statistical software (version 4.4.3, R Core Team, 2025) using the RStudio environment.

Considering the normality distribution of the data (Shapiro-Wilk test; P > 0.09 for all variables), paired t-test was used to compare the MVC peak torque, the length of the muscle and the coefficient of variation of torque for both hip positions.

For the motor unit analysis, linear mixed-effect models (LMMs) were applied to compare the mean discharge rate (trapezoidal and ramp-up tasks), CoV-ISI (trapezoidal task), recruitment threshold (ramp-up task), and discharge rate at recruitment (ramp-up task) between hip positions. This statistical model was used as it accounts for the non-independence of data points within each participant, which is particularly useful given the hierarchical nature of motor unit data[53]. To ensure statistical validity we also visually inspected the residuals quantile-quantile plot (Q-Q plot) for each model, and, for all cases, it followed a normal pattern of distribution. Two random-intercept models separated by force level were used, with hip angle (90° and 180°) and muscle (VM and VL) as fixed effects, and participant as the random effect. For the ramp-up contractions, the same random-intercept model was used. For the RF muscle, separate models were used as only non-matched motor units were included in this analysis. LMMs were applied using the package *lmerTest* [54] and the Kenward-Roger method was used to estimate the degrees of freedom and *P*-values. The package *emmeans* was used for multiple comparisons and to estimate marginal means with 95% confidence intervals.

To test if the proportion of change of the peak twitch at low and high frequency were significantly different from 0 a one-sample t-test was used separately for each frequency of stimulation. In addition, to test if the proportions of change were different between lower and higher frequencies they were compared using a paired t-test. In both cases, t-tests were used as the data was normally distributed (Shapiro-Wilk test; P>0.05).

For all statistical comparisons, the statistical significance was set at an α of 0.05. The effect sizes for all results were calculated using the *eff_size* function, which estimates the Cohen’s d from the outputs of the *emmeans* package (small > 0.2, moderate > 0.5, and large > 0.8) [55]. When needed for the paired t-test, the effect size was corrected for correlation. All individual data of motor unit discharge times recorded at hip 90° and hip 180° are available at https://doi.org/10.6084/m9.figshare.31236067.

## Results

While the RF muscle length significantly increased from 90° to 180° (hip at 90°: 83.24% ± 3.58; hip at 180°: 92.66% ± 5.06; t(10) = 7.739, *P* < 0.001, Cohen’s d = 2.057), the VL (hip at 90°: 87.41% ± 5.7; hip at 180°: 87.23% ± 5.13%; t(10) = 0.247, *P* = 0.810, Cohen’s d = 0.031) and VM (hip at 90°: 82.35 ± 7.54%; hip at 180° 83.51% ± 7.31%; t(10) = 2.032, *P* = 0.07, Cohen’s d = 0.155) muscle lengths did not significantly change between hip positions. To assess the effects of changing RF length on isometric knee extension torque production, the MVC peak torque and the torque steadiness during trapezoidal contractions (10% and 30% MVC) between hip positions (90° and 180°) were compared. No significant differences were found in MVC peak torque between 90° and 180° hip angles (hip at 90°: 188.3 ± 84.5 Nm; hip at 180°: 176 ± 73 Nm; paired t-test, t(17) = 1.438, *P* = 0.169, Cohen’s d = 0.149). Similarly, no significant differences in the coefficient of variation of torque were found between hip positions for both 10% MVC (hip at 90°: 4.40 ± 1.61%; hip at 180°: 4.11 ± 1.11%; paired t-test, t(16) = 0.822, *P* = 0.423, Cohen’s d = 0.200) and 30% MVC (hip at 90°: 4.64 ± 1.96%; hip at 180°: 4.20 ± 2.02%; t(15) = 0.672, *P* = 0.511, Cohen’s d = 0.220).

### Motor unit discharge behavior with changes in RF length (Experiment 1)

To evaluate whether the motor unit discharge properties (i.e., mean discharge rate and COV-ISI) were affected by changes in RF length, HDsEMG signals were decomposed into motor unit spike trains. The number of motor units per muscle, force level and hip position are provided in Supplementary Material 1. Briefly, the number of motor units tracked across hip positions at 10% MVC and 30% MVC was 3 ± 2 (VM) and 7 ± 8 (VL), and 2 ± 2 (VM) and 5 ± 8 (VL), respectively. The average number of RF motor units decomposed at 10% and 30% MVC for the hip at 90° was 4 ± 3 and 2 ± 1, respectively, and for the hip 180° was 3 ± 2 and 3 ± 1. The average silhouette value for these motor units was 0.92 ± 0.03 for the VM, 0.93 ± 0.02 for the VL and 0.91 ± 0.03 for the RF.

At 10% MVC, the mean motor unit discharge rate was higher at 180° compared to 90° hip angle (LMM, main effect of hip angle: F = 5.565, *P* = 0.019, Cohen’s d = 0.34; **Figure 4A**), with no significant interaction between hip angle and muscle (LMM, interaction effect: F = 0.154, *P* = 0.635; Cohen’s d range [0.014 - 0.423]). In contrast, at 30% MVC, no significant differences in mean discharge rate was observed between hip positions (LMM, main effect of hip angle: F = 1.261, *P* = 0.263; Cohen’s d = 0.21; **Figure 4B**), and also no significant interaction between hip angle and muscle (LMM, interaction effect: F = 1.2385, *P=*0.267, Cohen’s d range [0.018 – 1.47]). For the non-matched RF motor units, the mean discharge rate was similar between hip positions at 10% MVC (LMM, main effect of hip angle, F= 0.0014, *P =* 0.97, Cohen’s d = 0.009) and 30% MVC (LMM, main effect of hip angle, F= 0.4422, *P* = 0.511, Cohen’s d = 0.237).

**Figure 4.**
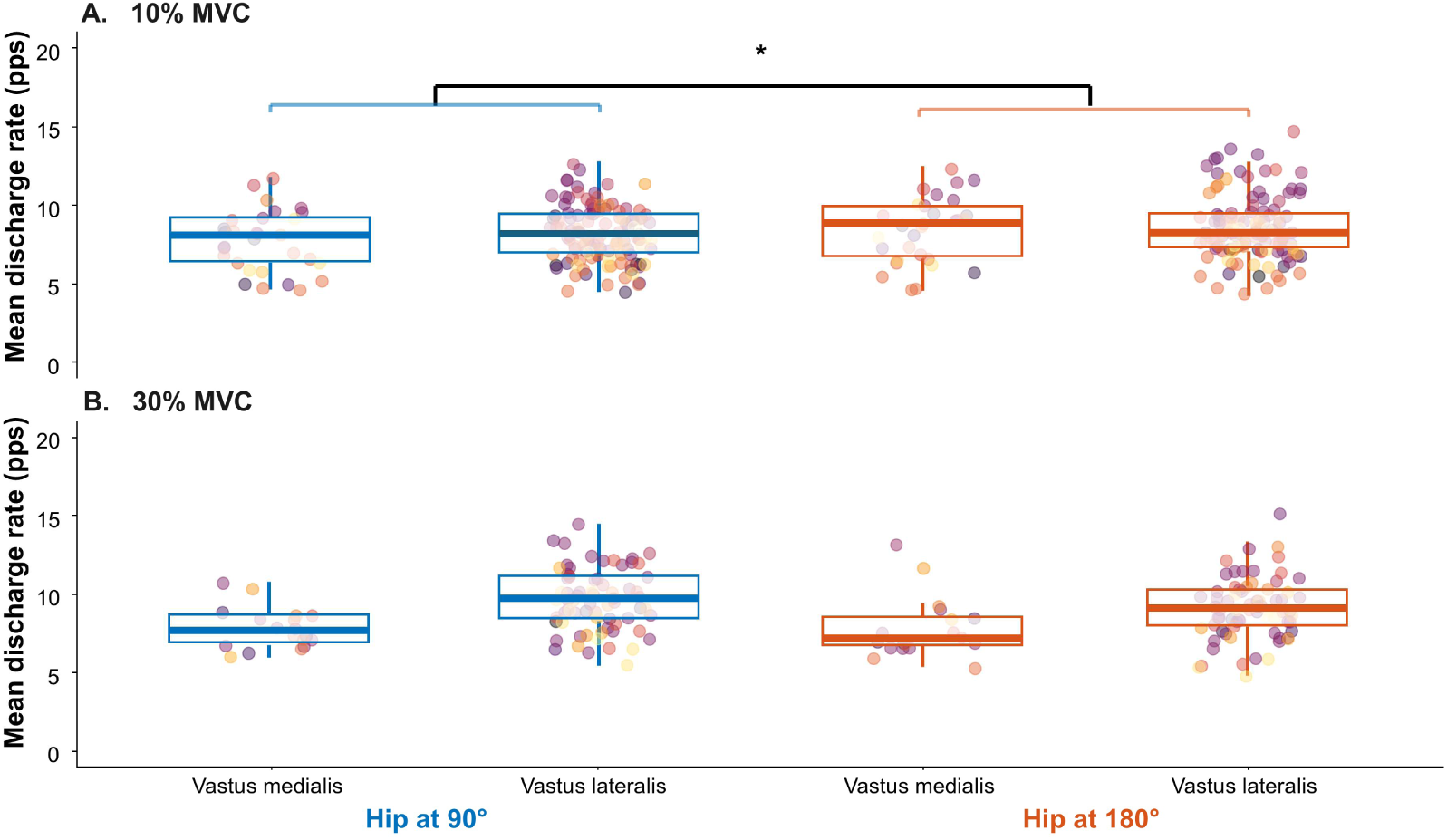
Mean discharge rate at the trapezoidal tasks for vasti muscles with the hip positioned at 90° (rectus femoris was shortened; blue) and at 180° (rectus femoris was lengthened; orange). Results are shown separately for 10% MVC (A) and 30% MVC (B). The central line of the boxplots indicates the median, the top and bottom line the first and third quartiles, and the two whiskers corresponds to the 1.5 interquartile range. Each circle represents individual motor unit results and each color represent one participant. * indicates P < 0.05.

For both VM and VL motor units, the CoV-ISI did not significantly differ between hip positions at 10% MVC (LMM, main effect of hip angle: F = 0.059, *P* = 0.808, Cohen’s d = 0.035) and 30% MVC (LMM, main effect of hip angle: F = 0.108, *P* = 0.743, Cohen’s d = 0.061). No significant effect of interaction between hip angle and muscle was observed at 10% MVC (LMM, interaction effect: F = 0.016, *P =* 0.898, Cohen’s d range [0.016 – 0.569]) neither at 30% MVC (LMM, interaction effect: F = 0.038, *P =* 0.844, Cohen’s d range [0.024 – 1.123]). For the non-matched RF motor units, the CoV-ISI was similar between hip positions at 10% MVC (LMM, main effect of hip angle, F= 3.184, *P =* 0.079, Cohen’s d = 0.466) and at 30% MVC (LMM, main effect of hip angle, F= 1.084, *P =* 0.306, Cohen’s d =0.363). The mean values of mean discharge rate and CoV-ISI, along with 95% confidence intervals, for matched units of VM and VL and non-matched of RF are reported in **Table 1**.

**Table 1:** Characteristics of motor unit discharge rate from vastus medialis, rectus femoris and vastus lateralis muscles separately per force level (10% and 30%) and hip angles (90° and 180°). Matched units reported for VM and VL, and non-matched units for the RF. Values are reported as estimated marginal means with lower and upper bound from 95% confidence level. Mean discharge rate is in pulses per second (pps) and coefficient of variation of interspike interval (CoV-ISI) in percentage.

| Muscle | MVC (%) | Angle (°) | Mean discharge rate (pps) | CoV- ISI (%) |
| --- | --- | --- | --- | --- |
| VM | 10 | 90 | 7.67 [7.06 - 8.29] | 16.10 [14.13 - 18.06] |
|  |  | 180 | 8.17 [7.56 - 8.79] | 16.18 [14.21 - 18.14] |
|  | 30 | 90 | 7.86 [7.06 - 8.66] | 19.89 [17.35 - 22.42] |
|  |  | 180 | 7.85 [7.06 - 8.65] | 20.37 [17.84 - 22.91] |
| VL | 10 | 90 | 8.15 [7.85 - 8.46] | 14.34 [13.37 - 15.32] |
|  |  | 180 | 8.49 [8.18 - 8.79] | 14.60 [13.62 - 15.57] |
|  | 30 | 90 | 10.27 [9.88 - 10.67] | 15.46 [14.19 - 16.72] |
|  |  | 180 | 9.62 [9.23 - 10.02] | 15.58 [14.31 - 16.85] |
| RF | 10 | 90 | 8.43 [7.86 - 9.00] | 22.73 [20.80 - 24.66] |
|  |  | 180 | 8.46 [7.76 - 9.15] | 19.84 [17.49 - 22.19] |
|  | 30 | 90 | 8.22 [7.34 - 9.10] | 22.29 [19.31 - 25.27] |
|  |  | 180 | 7.90 [7.02 - 8.78] | 23.95 [20.97 - 26.93] |

### Ramp up contractions (Experiment 2)

For the ramp-up task, 2 ± 1 motor units were tracked for the VM and 6 ± 6 motor units for the VL. For both muscles, motor unit recruitment threshold significantly increased from 14.7 [12.9 – 16.5] % MVC with the hip at 90° to 16.6 [14.8 – 18.4] % MVC with the hip at 180° (LMM main effect of angle, F = 5.728, *P =* 0.017, Cohen’s d = 0.407), with no interaction effect between hip angle and muscle (LMM, interaction effect: F = 0.017, *P =* 0.896, Cohen’s d range [0.209 – 1.024]). To further explore whether differences in recruitment threshold could explain force-dependent changes observed in *Experiment 1*, motor units were divided into those recruited below 20% MVC and those recruited at or above 20% MVC. While motor units with higher recruitment thresholds were not significantly different for discharge rate at recruitment (hip at 90°: 7.73 [6.53 – 8.92] pps; hip at 180°: 7.21 [6.28 – 8.14] pps; LMM, F = 0.718, *P =* 0.403, Cohen’s d = 0.326; top panel in **Figure 5A**), motor units with lower recruitment thresholds increased their discharge rate at recruitment from 7.0[6.19 – 7.91] at 90° to 7.55 [6.77 – 8.34] pps at 180° (LMM, main effect of hip angle: F = 4.631, *P =* 0.032, Cohen’s d = 0.449; bottom panel in **Figure 5A**). Conversely, no significant differences in mean discharge rate were observed between hip positions for both higher recruitment threshold (hip at 90°: 8.6 [7.68 – 9.52] pps; hip at 180°: 8.29 [7.52– 9.06]; LMM, main effect of hip angle: F = 0.654, *P* = 0.426, Cohen’s d = 0.321) and lower recruitment threshold (hip at 90°: 9.29 [8.50 – 10.1] pps; hip at 180°: 9.57 [8.75 – 10.4] pps; LMM, main effect of hip angle: F = 1.063, *P* = 0.3037, Cohen’s d = 0.215) motor units.

**Figure 5.**
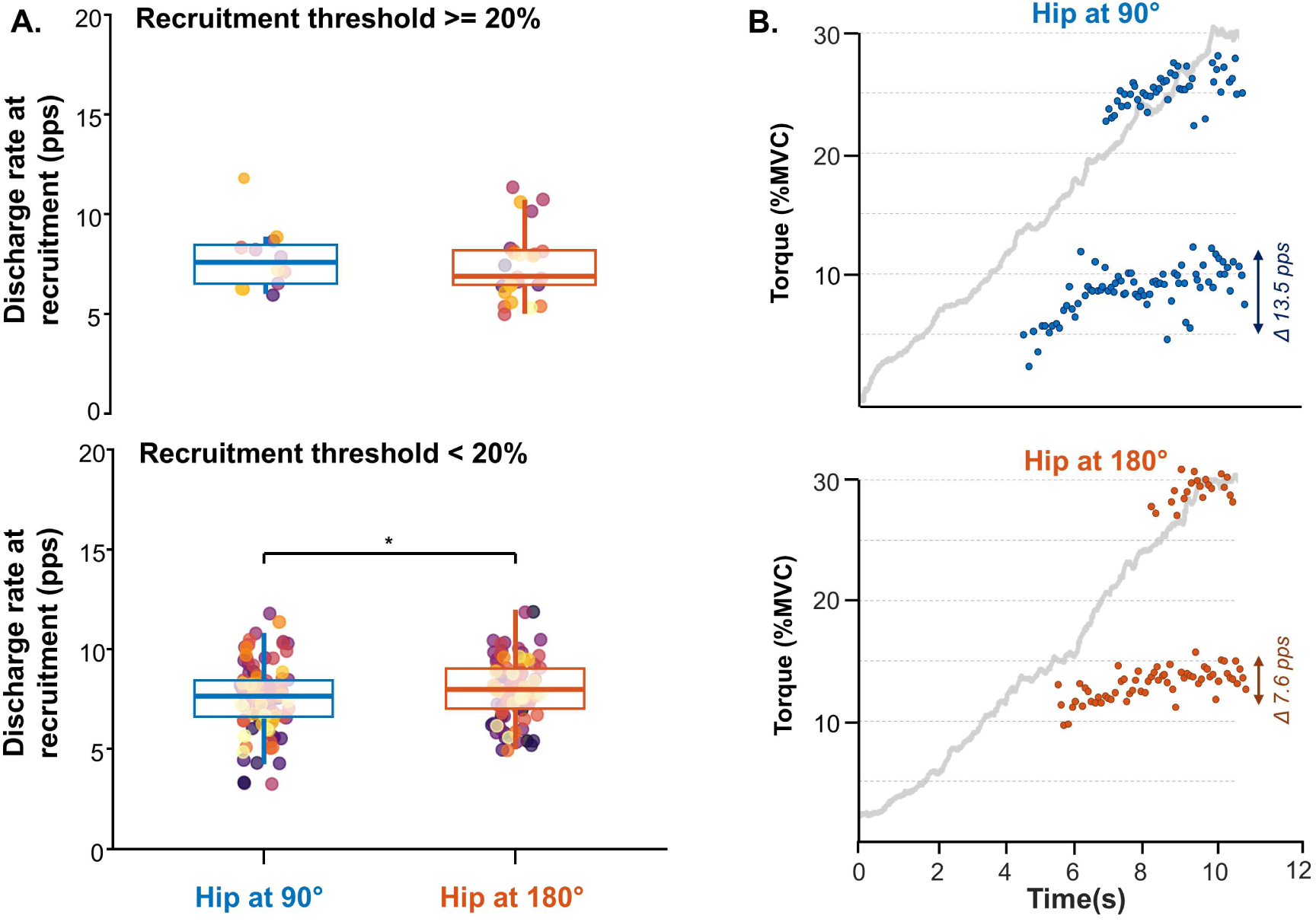
Ramp results with the hip positioned at 90° (rectus femoris was shortened; blue) and at 180° (rectus femoris was lengthened; orange). (A) Group results for the discharge rate at the recruitment between hip at 90° and 180°. Top panel corresponds to the discharge rate at recruitment for the units recruited at or above 20% MVC and bottom panel to the units recruitment below 20% MVC. The central line of the boxplot corresponds to the median value, the top and bottom lines the first and third quartiles, and the whiskers the 1.5 interquartile range. Each circle corresponds to a motor unit and each color to a participant. * Indicates P < 0.05. (B) Representative case showing instantaneous discharge rates of two motor units, one with lower and one with higher recruitment thresholds, during the ramp up contractions with the hip at 90° (top) and 180° (bottom).

**Figure 5B** shows a representative example of the ramp-up knee extension task for both 90° (top) and 180° (bottom) hip positions. In agreement with the group results, there is an increase in the motor unit recruitment threshold from the top (hip at 90°) to the bottom panel (hip at 180°). Additionally, the unit recruited below 20% presented an increase in discharge rate at recruitment, while the unit recruited above 20% did not.

Regarding the RF, the number of non-matched units at 90° and 180° hip position was to 3 ± 1 and 2 ± 1, respectively. There was no difference between angles for the recruitment threshold (LMM, F = 0.960, *P* = 0.331, Cohen’s d = 0.250). In addition, no significant differences regarding discharge rate at recruitment and mean discharge rate were observed for both lower recruitment threshold (discharge rate at recruitment: LMM, F = 0.343, *P* = 0.562, Cohen’s d = 0.208; mean discharge rate: LMM, F = 0.343, *P* = 0.343, Cohen’s d = 0.311) and higher recruitment threshold units (discharge rate at recruitment: LMM, F = 2.341, *P* = 0.140, Cohen’s d = 0.935; mean discharge rate: LMM, F = 2.341, *P* = 0.140, Cohen’s d = 0.591).

### Evoked knee extension torque twitches (Experiment 3)

To assess the effect of changes in RF muscle length from evoked twitches on the vasti muscles, a second set of experiments was conducted in a subgroup of seven participants where electrical stimuli were applied simultaneously over the VM and VL for the hip positioned at 90° and 180°. The element-wise average twitches for the six participants are displayed in **Figure 6** for both frequency stimulations at the same amplitude of 15 mA. It is possible to observe a reduction in the average twitch between 90° (blue line) and 180° (orange line) hip angles for 10 Hz stimulation (**Figure 6A**), but not for 20 Hz stimulation (**Figure 6B**). In agreement, the group results revealed that the proportion of peak twitch change was significantly lower than 0 at 10 Hz (−45.81 ± 43.71%; One sample t-test, t(6)=2.773, *P* = 0.032, Cohen’s d = 1.048), while no significant changes were observed at 20 Hz (7.86 ± 79.91; t(6)=0.260, *P* = 0.803, Cohen’s d = 0.098). In addition, the lower frequency presented a significantly greater proportion of change than the higher frequency stimulation (t(6) = 3.215, *P* = 0.018, Cohen’s d = 0.520).

**Figure 6.**
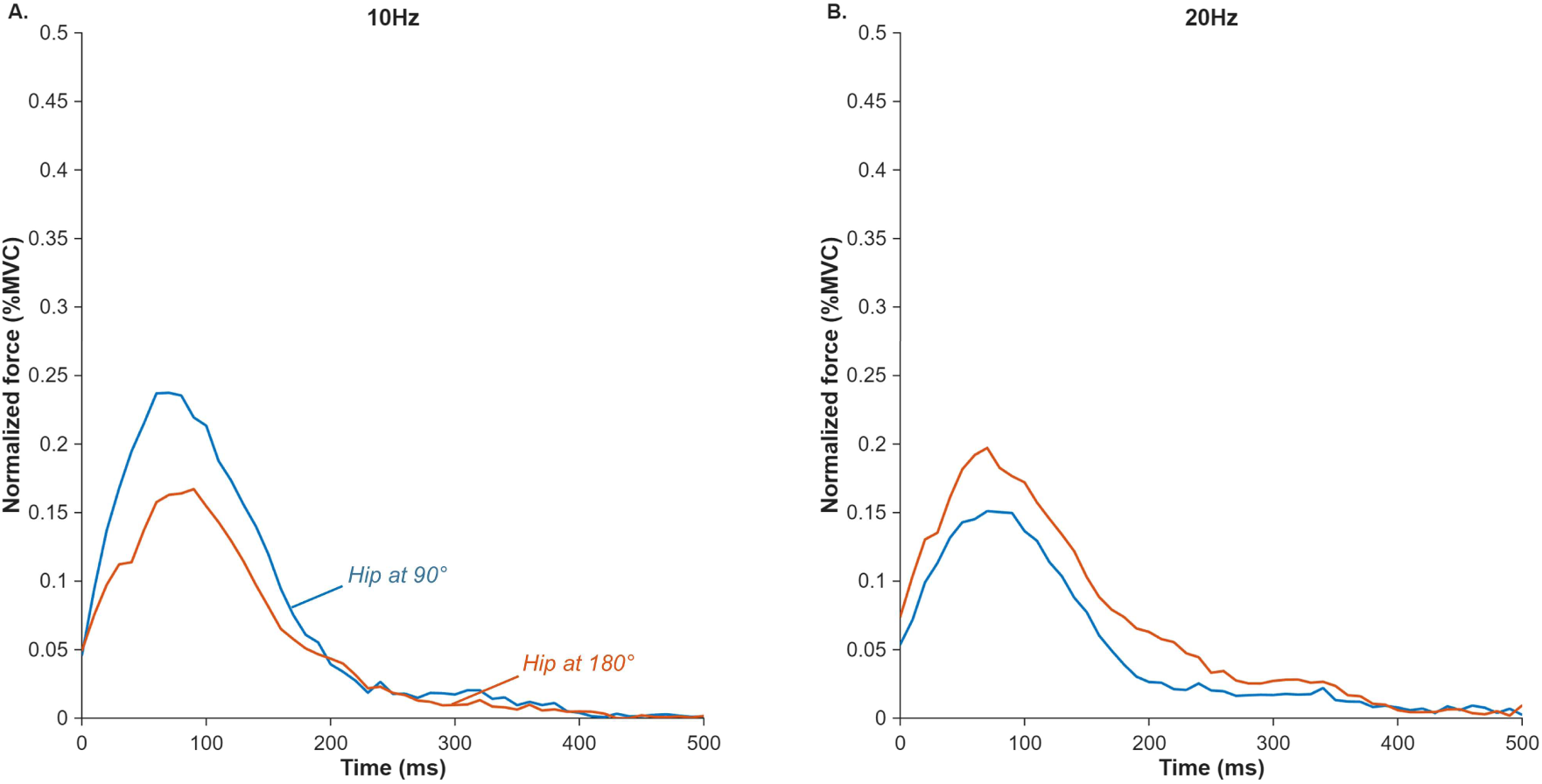
Averaged muscle twitch across participants for the hip angle at 90° (rectus femoris was shortened; blue) and at 180° (rectus femoris was lengthened; orange). The average twitch profile representing the combined contribution of the vasti muscles was estimated using a deconvolution method. (A) corresponds to the stimulation frequency of 10 Hz and (B) corresponds to stimulation frequency of 20 Hz.

## Discussion

The present study investigated how changes in the length of the biarticular RF muscle influence both central and peripheral motor unit properties of its synergistic muscles (VM and VL). The primary findings showed that lengthening the RF, achieved by changing the hip joint angle from 90° to 180° and confirmed by ultrasound measurements, led to increased mean discharge rates of VM and VL motor units at 10% MVC, but not at 30% MVC. Two additional experiments were conducted to further explore these force-dependent changes. In *Experiment 2*, increases in recruitment threshold and discharge rate at recruitment were observed in the lengthened RF condition, but only for motor units recruited below 20% MVC. In *Experiment 3*, conducted in a subgroup of participants, electrically evoked twitch responses of the vasti muscles were significantly reduced at 10 Hz but not 20 Hz when the hip was positioned at 180°. Taken together, these findings suggest that, even though alterations in RF length did not change its motor unit properties, it affects both central and peripheral motor unit properties of VM and VL. Notably, as discussed below, force-dependent changes observed in motor unit discharge rate appear to be closely linked to alterations in the contractile properties of the vasti muscles resulting from changes in RF length.

Muscle force production and control rely on the interaction between the effective neural drive to the muscle (i.e., the cumulative spike train of active motor units) and the mechanical properties of the muscle-tendon unit [1,10]. To examine this relationship, several studies have investigated how alterations in muscle length influence the neural drive, either through direct assessments of motor unit discharge behavior [28–32] or indirectly via surface EMG amplitude [43,56]. For instance, altering knee joint position from extension to flexion has been shown to increase recruitment thresholds and decrease the mean discharge rates of gastrocnemii motor units during both dynamic [31] and isometric [32,57] plantar flexion tasks. Similarly, a reduction in voluntary activation has been reported in the rectus femoris when moving from a seated to a supine position [17]. These results suggest that altering the muscle length can directly influence the discharge and activation behavior of the muscle itself. However, this is not a consensus in the literature [28–32,57]. During dorsiflexion contractions, for example, motor unit discharge rates remained unchanged across different lengths of tibialis anterior muscle [28,29,33]. In agreement with this, this study also showed that alterations in the RF length did not alter its mean discharge rate. Collectively, these findings suggest that mean discharge rate variations with muscle length may be muscle dependent.

In the present study, we also examined the effect of changes in the length of the RF muscle-tendon unit on the motor unit behavior of its synergistic counterparts, VM and VL. At the low force level, mean discharge rates of vasti motor units increased when the RF was lengthened. Importantly, because no significant differences were found in MVC or force steadiness between hip positions, the increased discharge rates in the vasti are unlikely to reflect changes in task demands. Instead, they likely represent a compensatory neural strategy to maintain knee extension torque in response to an altered mechanical contribution of the lengthened RF at a hip angle of 180°. This interpretation is consistent with previous observations in the triceps surae, where changes in the gastrocnemius muscle length were associated with altered motor unit discharge behavior in its synergist soleus muscle. Several mechanisms may underlie this neural-mechanical interaction between a muscle and its synergists. Although active force is generated longitudinally along muscle fibers according to their architecture, total muscle force (active and passive) is transmitted not only along the line of action but also transversely to adjacent muscles through connective tissue linkages [58]. As a result, force produced by the RF may influence the mechanical environment of the VM and VL via transverse force transmission, potentially contributing to the increased discharge rates observed in the vasti when RF length was altered. Although RF was the only muscle whose length was manipulated, confirmed by ultrasound measurements,^58^ the increased vasti motor unit discharge rates at 180° may also reflect changes in afferent feedback. In particular, heteronymous Ia afferent projections from the RF to synergistic muscles such as the VM and VL may contribute, as these pathways are known to synapse onto motoneurons of adjacent synergistic muscles [59–61]. This interpretation aligns with previous findings indicating that the VM and VL share a substantial portion of their synaptic inputs, consistent with their joint contribution to knee extension and stabilization of the patellofemoral joint [36,62].

Interestingly, the increase in mean discharge rate of vasti motor units induced by RF lengthening was observed only at 10% MVC and not at 30% MVC. Although the precise physiological mechanisms underlying this force-dependent effect are difficult to isolate, several factors may account for this finding. One possibility could be that changes in RF muscle length modulate vasti motor unit discharge behavior through recruitment-threshold-dependent mechanisms. Previous studies have shown that low- and high-threshold motor units adjust their discharge rates differently under several conditions, including fatigue, muscle pain, and changes in muscle length during different types of contraction [31,32,57,63–65]. To test whether such mechanism could explain our observations, we performed ramp-up contractions up to 30% MVC and analyzed motor unit recruitment thresholds and discharge rates (*Experiment 2*). When the RF was lengthened, vasti motor units exhibited higher discharge rates at recruitment, but only for units recruited below 20% MVC. These findings extend current knowledge by indicating that changes in muscle length at low force levels influence synergistic muscle behavior primarily through modulation of low-threshold motor unit discharge rates. ^63^

In addition to neural mechanisms, force dependent changes may also be explained by musculotendinous mechanical properties, particularly the behavior of passive elastic components. One plausible explanation relates to the nonlinear stress–strain and viscoelastic properties of the tendon. When the RF is elongated (hip joint at 180°), at the onset of contraction its tendon may already be positioned beyond the toe region of the stress–strain curve[66,67]. However, at low force levels (10% MVC), the generated tension may still be insufficient to effectively engage and sustain force transmission, thereby requiring increased neural activation, as reflected by the increased vasti discharge rates. As force level increases to 30% MVC, the higher tension may improve mechanical efficiency of force transmission [68], reducing the need for compensatory increases in motor unit discharge rate.^69^ To further explore this interpretation, we examined whether force-dependent changes were associated with alterations in vasti twitch contractile properties. *Experiment 3* revealed significantly reduced peak twitch amplitude at a hip angle of 180° compared to 90° (**Figure 6**), but only at low stimulation frequencies, with no differences observed at higher frequencies. Given that muscle force output results from the convolution of motor unit spike trains with the average twitch response of the muscle [4,69], a reduction in twitch amplitude would necessitate higher motor unit discharge rates to maintain the required force output, which is consistent with our findings. Together, these results support the idea that at low force levels, greater neural input from the vasti is required to compensate for altered mechanical behavior associated with RF lengthening. Thus, both diminished contractile responses and altered tendon tension likely contribute to the increased vasti motor unit discharge rates observed at 10% MVC, but not at 30% MVC.

A final consideration concerns the lack of significant changes in MVC between hip positions, which aligns with some previous findings [56,70], but contrasts with other studies reporting reduced voluntary isometric knee extension torque under similar protocols [17–19]. This discrepancy may be explained by evidence suggesting that the knee angle has a more pronounced effect on quadriceps torque production than hip angle [18,71]. Additionally, when analyzing the contribution of each muscle to total quadriceps torque, the RF contributes the least[72], which could also explain the lack of difference between hip positions. Since the primary goal of this study was to isolate the effects of RF lengthening, the knee angle was kept constant across conditions.

Some limitations of this study should be noted. The relatively low yield of decomposed motor units from the RF muscle may have limited statistical power. RF is a challenging muscle for HDsEMG decomposition [6], so inferences should be made cautiously. Moreover, we were unable to match motor units of the RF muscle itself. Because the protocol involved altering the RF length, the placement of the electrode grid was not sufficiently consistent across hip positions to allow reliable comparisons. In addition, we could not collect data from vastus intermedius, but since its contribution to the total torque is similar to the other vasti [72], we believe that motor unit behavior of this muscle would follow a similar pattern to the one reported in this study for the VM and VL.

In conclusion, this study provides insights into how mechanical and neural characteristics of synergistic monoarticular muscles adapt after changing the length of one biarticular muscle during an isometric task. Here we focused on the behavior of quadriceps muscles, especially VM and VL, in response to changes in length of the biarticular RF muscle. Our findings indicated that manipulating the hip position can influence the neural drive of vasti motor units in a force-dependent manner. From a practical perspective, these results suggest that hip position should be considered when designing strength training or rehabilitation programs, as altering RF length may indirectly modulate the activation strategies of its synergistic muscles. In addition to changing the hip angle, other interventions, such as static stretching, could be explored to assess potential changes in the control of synergistic muscles [73,74].

## Author contributions

MAS, HVC, JGI & FN designed the study. MAS, CC, EP & JGI acquired the data. MAS, HVC & FN analyzed the data. MAS, HVC & FN wrote the manuscript. HVC, JGI, LFO & FN provided critical revision. All authors interpreted the data, contributed to the manuscript, reviewed it, approved the content of the final version, and agree to be accountable for all aspects of the work. All persons designated as authors qualify for authorship, and all those who qualify for authorship are listed.

## Acknowledgments

Funded by the European Union. Views and opinions expressed are, however, those of the author(s) only and do not necessarily reflect those of the European Union or the European Research Council Executive Agency. Neither the European Union nor the granting authority can be held responsible for them. This study was funded by the European Research Council Consolidator Grant INcEPTION contract n. 101045605 (to FN). J Greig Inglis was supported by the Marie Skłodowska-Curie Actions Grant ‘MUDecomp’, agreement no. 101151712. Liliam F. Oliveira was supported by the Fundação Carlos Chagas Filho de Amparo à Pesquisa do Estado do Rio de Janeiro (FAPERJ), grant number E-26/210.128/2023.

## Data availability statement

All individual data of motor unit discharge times recorded at hip 90° and hip 180° are available at https://doi.org/10.6084/m9.figshare.31236067.

