## Supplementary material 1 for "Neural and mechanical properties of vastus lateralis and vastus medialis at different rectus femoris muscle lengths"

Table 1. Average motor unit yield considering matched and non-matched of the three muscles VM, VL and RF. Numbers are provided separately per task, force level and hip position.

| Muscle | Force level | Task | Hip position | Matching type | Average ± Standard deviation |
|---|---|---|---|---|---|
| VL | 10 | Trapezoidal | H90 | matched | 7±8 |
| VM | 10 | Trapezoidal | H90 | matched | 3±2 |
| VL | 30 | Trapezoidal | H90 | matched | 5±8 |
| VM | 30 | Trapezoidal | H90 | matched | 2±2 |
| VL | 30 | Ramp | H90 | matched | 6±6 |
| VM | 30 | Ramp | H90 | matched | 2±1 |
| VL | 10 | Trapezoidal | H180 | matched | 7±8 |
| VM | 10 | Trapezoidal | H180 | matched | 3±2 |
| VL | 30 | Trapezoidal | H180 | matched | 5±8 |
| VM | 30 | Trapezoidal | H180 | matched | 2±2 |
| VL | 30 | Ramp | H180 | matched | 6±6 |
| VM | 30 | Ramp | H180 | matched | 2±1 |
| RF | 10 | Trapezoidal | H90 | non-matched | 4±3 |
| VL | 10 | Trapezoidal | H90 | non-matched | 13±8 |
| VM | 10 | Trapezoidal | H90 | non-matched | 3±2 |
| RF | 30 | Trapezoidal | H90 | non-matched | 2±1 |
| VL | 30 | Trapezoidal | H90 | non-matched | 12±8 |
| VM | 30 | Trapezoidal | H90 | non-matched | 3±2 |
| RF | 30 | Ramp | H90 | non-matched | 3±1 |
| VL | 30 | Ramp | H90 | non-matched | 12±6 |
| VM | 30 | Ramp | H90 | non-matched | 3±2 |
| RF | 10 | Trapezoidal | H180 | non-matched | 3±2 |
| VL | 10 | Trapezoidal | H180 | non-matched | 16±11 |
| VM | 10 | Trapezoidal | H180 | non-matched | 3±2 |
| RF | 30 | Trapezoidal | H180 | non-matched | 3±1 |
| VL | 30 | Trapezoidal | H180 | non-matched | 16±10 |
| VM | 30 | Trapezoidal | H180 | non-matched | 3±2 |
| RF | 30 | Ramp | H180 | non-matched | 2±1 |
| VL | 30 | Ramp | H180 | non-matched | 14±7 |
| VM | 30 | Ramp | H180 | non-matched | 3±1 |
